# The loader complex Scc2/4 forms co-condensates with DNA as loading sites for cohesin

**DOI:** 10.1101/2021.12.14.472603

**Authors:** Sarah Zernia, Dieter Kamp, Johannes Stigler

## Abstract

The genome is organized by diverse packaging mechanisms like nucleosome formation, loop extrusion and phase separation, which all compact DNA in a dynamic manner. Phase separation additionally drives protein recruitment to condensed DNA sites and thus regulates gene transcription. The cohesin complex is a key player in chromosomal organization that extrudes loops to connect distant regions of the genome and ensures sister chromatid cohesion after S-phase. For stable loading onto the DNA and for activation, cohesin requires the loading complex Scc2/4. As the precise loading mechanism remains unclear, we investigated whether phase separation might be the initializer of the cohesin recruitment process. We found that, in absence of cohesin, budding yeast Scc2/4 forms phase separated co-condensates with DNA, which comprise liquid-like properties shown by droplet shape, fusion ability and reversibility. We reveal in DNA curtain and optical tweezer experiments that these condensates are built by DNA bridging and bending through Scc2/4. Importantly, Scc2/4-mediated condensates recruit cohesin efficiently and increase the stability of the cohesin complex. We conclude that phase separation properties of Scc2/4 enhance cohesin loading by molecular crowding, which might then provide a starting point for the recruitment of additional factors and proteins.

## Introduction

In eukaryotic interphase cells, DNA is folded and condensed to fit into the confines of the nucleus. This folding needs to be dynamic to allow biochemical reactions like DNA replication, repair, recombination or transcription to occur. DNA is folded into nucleosomes forming a beads-on-a-string-like structure, which compacts DNA around sevenfold (Oudet et al., 1975). Different histone modifications furthermore regulate accessibility of the DNA leading to the less densely packed gene-rich euchromatin and the more densely packed transcriptional inactive heterochromatin (Y. Kim & Yu, 2020). Additionally, the chromatin fiber is folded by loop formation into topologically associated domains (TADs), which exhibit high self-interaction and bring far-distance regions of the genome into close proximity (Dixon et al., 2012; Fudenberg et al., 2016; Pombo & Dillon, 2015). This formation of spatially organized structures promotes contacts between enhancers and promoters and is tightly linked to gene regulation (Pope et al., 2014; Szabo et al., 2019).

With its apparent ability to extrude loops of DNA, the cohesin complex is one of the key proteins in TAD formation (Davidson & Peters, 2021; Fudenberg et al., 2017). Furthermore, cohesin establishes sister chromatid cohesion (Guacci et al., 1997; Michaelis et al., 1997) most likely by simultaneously entrapping two DNA strands (Haering et al., 2008). To stably load onto DNA, cohesin requires a loader complex, called Scc2/4 in budding yeast (Ciosk et al., 2000; Collier et al., 2020; Murayama & Uhlmann, 2014). This heterodimer is shaped like a hook, provides two interaction sites for DNA (Kurokawa & Murayama, 2020) and can clamp cohesin on DNA by interfacing with all cohesin subunits (Chao et al., 2015; Higashi et al., 2020), which leads to increased lifetime of cohesin on DNA (Gutierrez-Escribano et al., 2019; Stigler et al., 2016). However, the cohesin-Scc2/4 interaction seems to be relatively transient as Scc2/4 was found to hop between different cohesin complexes (Rhodes et al., 2017) and is counteracted by Pds5 (Kikuchi et al., 2016; Petela et al., 2018) as well as Wapl (Haarhuis et al., 2017). Besides simple DNA clamping, Scc2/4 can stimulate the ATPase function of cohesin’s head domains leading to ATP hydrolysis, which is necessary for DNA entrapment inside the cohesin ring (Çamdere et al., 2015; Murayama & Uhlmann, 2014; Petela et al., 2018). Recently, it was found that Scc2/4 can bind to different sites of the cohesin ring, dependent on ATP, suggesting a two-step loading mechanism initialized by ATP-independent binding at the hinge region followed by a transition towards the ATPase head domains (Bauer et al., 2021). However, it remains unclear what initializes and drives the loading process.

Real-time single-molecule experiments have revealed that cohesin forms both DNA loops (Davidson et al., 2019; Y. Kim et al., 2019) and trans-tethers of DNA (Gutierrez-Escribano et al., 2019), exclusively in the presence of Scc2/4. Additionally, these experiments have shown that cohesin can unspecifically compact DNA (Gutierrez-Escribano et al., 2019), even across nucleosomes (Y. Kim et al., 2019), which is caused by the formation of large protein-DNA-co-condensates with liquid-like properties (Ryu et al., 2021). Phase separation was shown before to be an important mechanism in genome organization as it forms regions of increased protein and DNA concentration to facilitate for example gene transcription activation or repression (Boehning et al., 2018; Boija et al., 2018; Kent et al., 2020; Larson et al., 2017; Sabari et al., 2018; Strom et al., 2017). Furthermore, other DNA-binding proteins like HP1α (Keenen et al., 2020) and FUS (Zuo et al., 2021) were previously found to compact DNA by phase separation in a comparable fashion.

The requirement of Scc2/4 for the formation of cohesin-DNA co-condensates raises the question whether phase separation is not only an essential process for DNA organization but could also contribute to the cohesin loading process. To our surprise, we found that Scc2/4 alone is able to phase separate on DNA curtains and can form microscopic co-condensates with DNA in a concentration-dependent manner. We characterized these condensates in terms of preconditions, stability, reversibility and dynamics using DNA curtains and optical tweezers and found that Scc2/4 serves as a DNA bridging factor. Finally, we document preferred cohesin loading at Scc2/4-DNA-co-condensates and higher cohesin stability when it is loaded at these sites, suggesting a loading mechanism primed by phase separation.

## Results

### The cohesin loader Scc2/4 connects and compacts DNA molecules

To visualize the effects of the cohesin loader Scc2/4 on DNA, we immobilized ∼48.5 kbp dsDNA molecules derived from bacteriophage lambda to a lipid bilayer in a flow chamber that allowed us to exchange buffer components (see **Fig. S1A**). The YOYO-1-stained DNA molecules could diffuse freely in the absence of buffer flow (**Fig. 1A**). Upon injection of 8 nM Scc2/4, we observed the slow formation of puncta on the surface (**Fig. 1A**) that increased in fluorescence and compacted in size over time (**Fig. S1B,C**). The formation was strictly dependent on the presence of Scc2/4. Intrigued by this, to our knowledge, previously undescribed coagulation of DNA into clustered condensates by Scc2/4, we decided to characterize the properties and consequences of this phenomenon in more detail.

**Figure 1.**
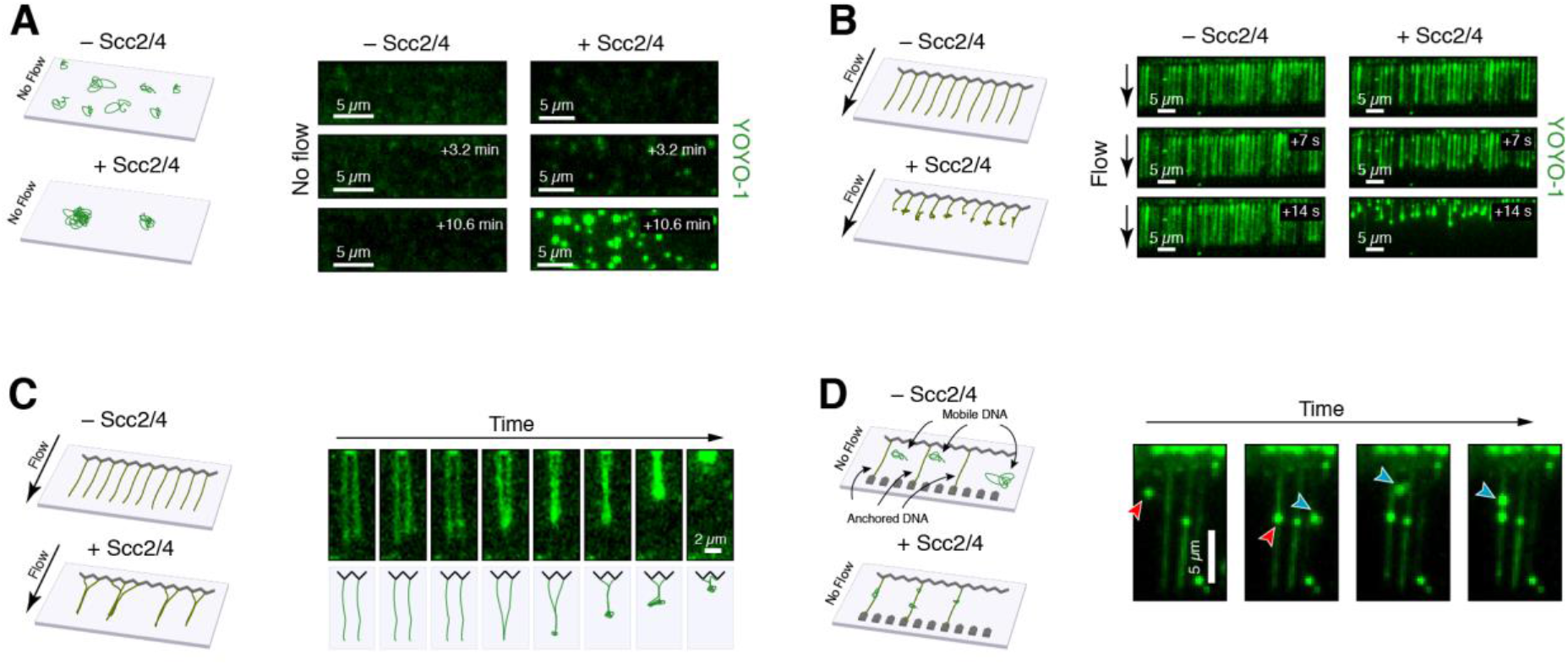
The cohesin loader Scc2/4 condenses and compacts DNA. (**A**) Diffusing DNA molecules, tethered to a lipid bilayer, form condensates after addition of 8 nM Scc2/4. (**B**) DNA curtains assay of flow-stretched DNA. Addition of 100 nM Scc2/4 leads to rapid compaction. (**C**) Braiding between neighboring flow-stretched DNA strands after Scc2/4 addition. (**D**) Mobile lipid-anchored DNA molecules (red and blue arrows) attach to double-tethered DNA molecules after Scc2/4 addition in the absence of flow.

Thus, we used single-tethered DNA curtains (Greene et al., 2010), where lipid-attached lambda DNA is flow-stretched across nanofabricated barriers allowing the investigation of multiple DNA molecules in an array format (**Fig. 1B**). While flow-stretched DNA remained at full length in the absence of Scc2/4, we observed rapid shortening and compaction of DNA upon the injection of 100 nM Scc2/4 (**Fig. 1B**). In a similar setting, we also observed Scc2/4-dependent braiding of adjacent DNA molecules, which typically started from the free end, followed by compaction (**Fig. 1C**), indicating that Scc2/4-mediated interactions led to both the coagulation of DNA molecules in trans as well as in cis. This protein-dependent formation of DNA clusters and tethering in trans was also apparent in double-tethered DNA curtains, where we observed an Scc2/4-dependent recruitment of mobile lipid-anchored DNA molecules (red and blue arrows in **Fig. 1D**) to stretched surface-anchored DNA molecules.

### Large-scale co-condensation of Scc2/4 with DNA occurs at physiological conditions

Co-condensation of nucleic acids with proteins has mainly been described and observed by the formation of macroscopic phase-separated droplets in solution (Keenen et al., 2020; Morin et al., 2020; Zhou et al., 2019; Zuo et al., 2021). To test whether Scc2/4-DNA co-condensates also form phase-separated droplets, we titrated varying concentrations of Scc2/4 to a 30 nM solution of DNA fragments of ∼2.9 kbp and checked for condensate formation by bright-field microscopy. Strikingly, we observed the formation of visible phase-separated droplets in an Scc2/4-concentration-dependent manner, with droplets appearing at concentration above 300 nM Scc2/4 (**Fig. 2A**). Note that condensates exclusively formed in presence of DNA. We therefore conclude that the observed clustering, coagulation and compaction of single DNA molecules is, in fact, co-condensation of Scc2/4 and DNA.

**Figure 2.**
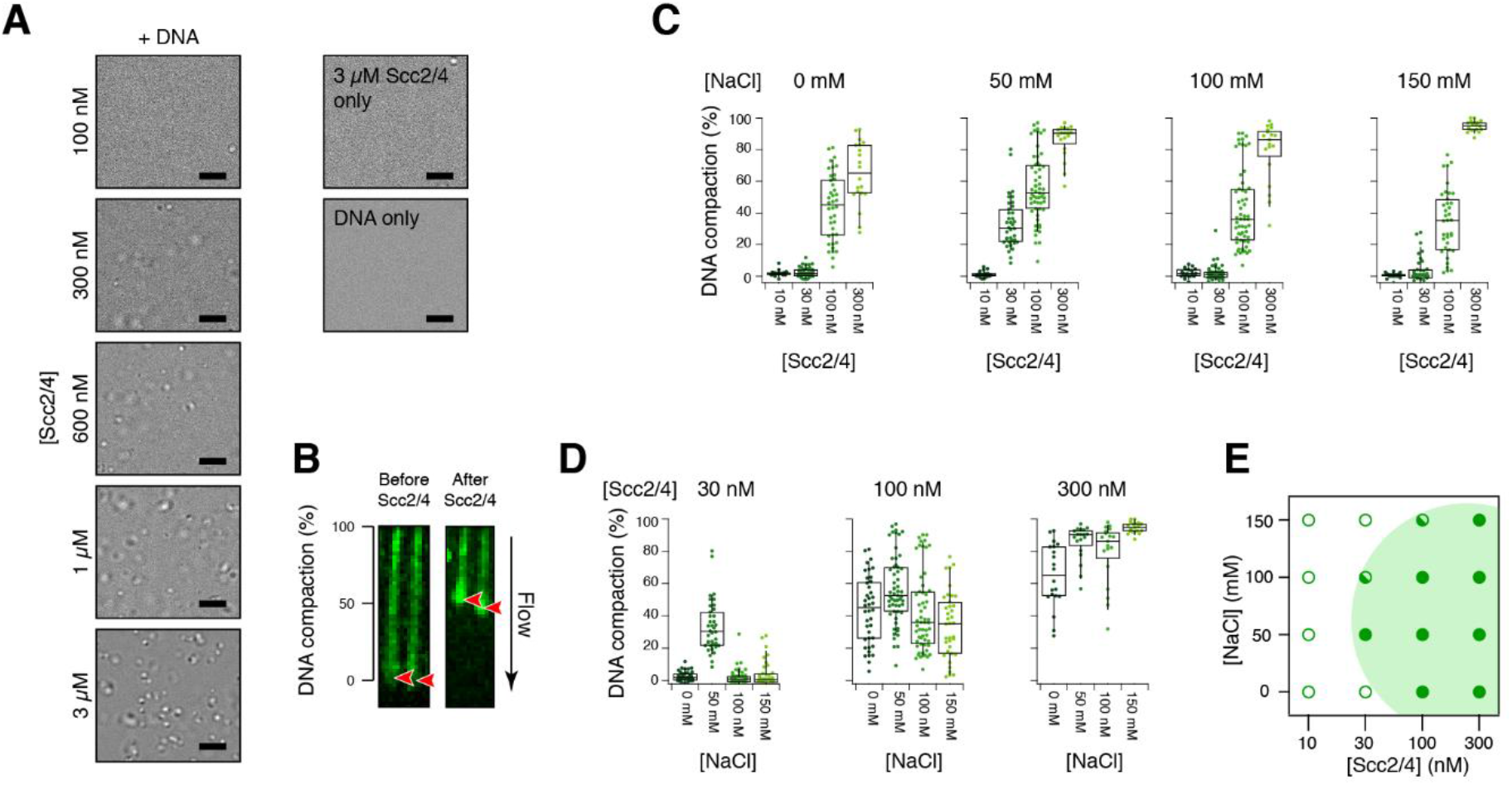
Quantification of DNA condensation from single molecule and bulk microscopy experiments. (**A**) Bright-field microscopy images of large co-condensates of Scc2/4 and 30 nM 2686 bp DNA fragment. Scale bar: 10 μm. (**B**) Example of DNA compaction by short pulses of Scc2/4. Red arrows show the free DNA end before and after compaction. (**C**) Compaction of single lambda DNA molecules after injection of 50 μl pulses of Scc2/4 at the indicated concentrations, at varying concentrations of NaCl. For 10 nM and 300 nM pof Scc2/4, 19 DNA strands were analyzed, for 30 nM and 100 nM, 38 DNA strands were analyzed. (**D**) Rearranged data of (**C**) showing an optimal salt concentration for condensation. (**E**) Phase diagram of DNA condensation from single molecule experiments. Open circles: no condensation; closed circles: condensation, half-filled circles: condensation occurred occasionally. The green area in the background represents the range where condensation was observed, to guide the eye.

To further explore the range of concentrations that led to the co-condensation of single DNA molecules, we returned to DNA curtain experiments. We injected 50 μl volumes of varying concentrations of Scc2/4 into a DNA curtain flow chamber with a flow rate of 0.1 ml/min, which exposed DNA to protein for ∼60 s (**Fig. S2A**). We then determined the degree of DNA compaction by measuring the DNA length before and after Scc2/4-exposure (**Fig. 2B**). The compaction was positively correlated with the concentration of Scc2/4 (see also **Fig. S2B** for examples of DNA compaction). While at 10 nM Scc2/4 the DNA remained essentially fully extended, 300 nM Scc2/4 compacted the DNA almost completely (**Fig. 2C**). We note that the reported Scc2/4 concentrations should be compared only within the context of the flow-stretched compaction assay. Lower concentrations still may induce condensation on relaxed DNA, albeit with a longer time constant (e.g. **Fig. 1A**). As protein-DNA co-condensation often relies on electrostatic interactions (Erdel & Rippe, 2018), we tested the influence of monovalent salt on DNA compaction by varying the NaCl concentration between 0 mM and 150 mM. However, while compaction was significantly enhanced at 50 mM NaCl, the overall dependency on salt concentration was weak (**Fig. 2C,D**). The data is summarized in the phase diagram of **Fig. 2E**, showing the range of Scc2/4 and NaCl concentrations where the formation of co-condensates was observed. Importantly, we observed significant compaction of DNA at near physiological salt (150 mM NaCl) and Scc2/4 concentrations (∼300 nM, (Ho et al., 2018), **Fig. 2C**, right), suggesting that Scc2/4-mediated DNA condensation may also occur in vivo.

### Scc2/4 serves as a DNA-bridging factor in condensates

Intrigued by the observation that the cohesin loader Scc2/4 can lead to the formation of DNA-DNA contacts on its own, without the help of the SMC complex cohesin, we next sought to characterize the mechanism of Scc2/4-mediated DNA condensate formation in greater detail. Condensation of biomolecules generally requires binding multivalency of the condensate components. Many phase separating proteins cluster by homo-interactions through intrinsically disordered domains (IDRs), which form a molten-globule-like state through hydrophobic interactions by excluding water molecules (LLPS, liquid-liquid phase separation, (Dignon et al., 2020; Erdel & Rippe, 2018; Razin & Ulianov, 2020)). However, prediction software does not indicate that there are extended IDRs in Scc2/4 (the unstructured N-terminus of Scc2 is structured upon binding of Scc4, **Fig. S3**). In agreement with this assessment, washes with 20 % 1,6-hexanediol, a strong disruptor of hydrophobic interactions that has been used extensively in studies of LLPS (Ladouceur et al., 2020; Ryu et al., 2021), was not able to decondense Scc2/4-DNA co-condensates (**Fig. 3A**). In contrast, disruption of electrostatic interactions, such as the injection of 2M NaCl readily dissolved Scc2/4-DNA co-condensates and returned the DNA back to its extended form (**Fig. 3B**). We conclude that the co-condensation mechanism between Scc2/4 and DNA is likely driven by electrostatic rather than hydrophobic interactions.

**Figure 3.**
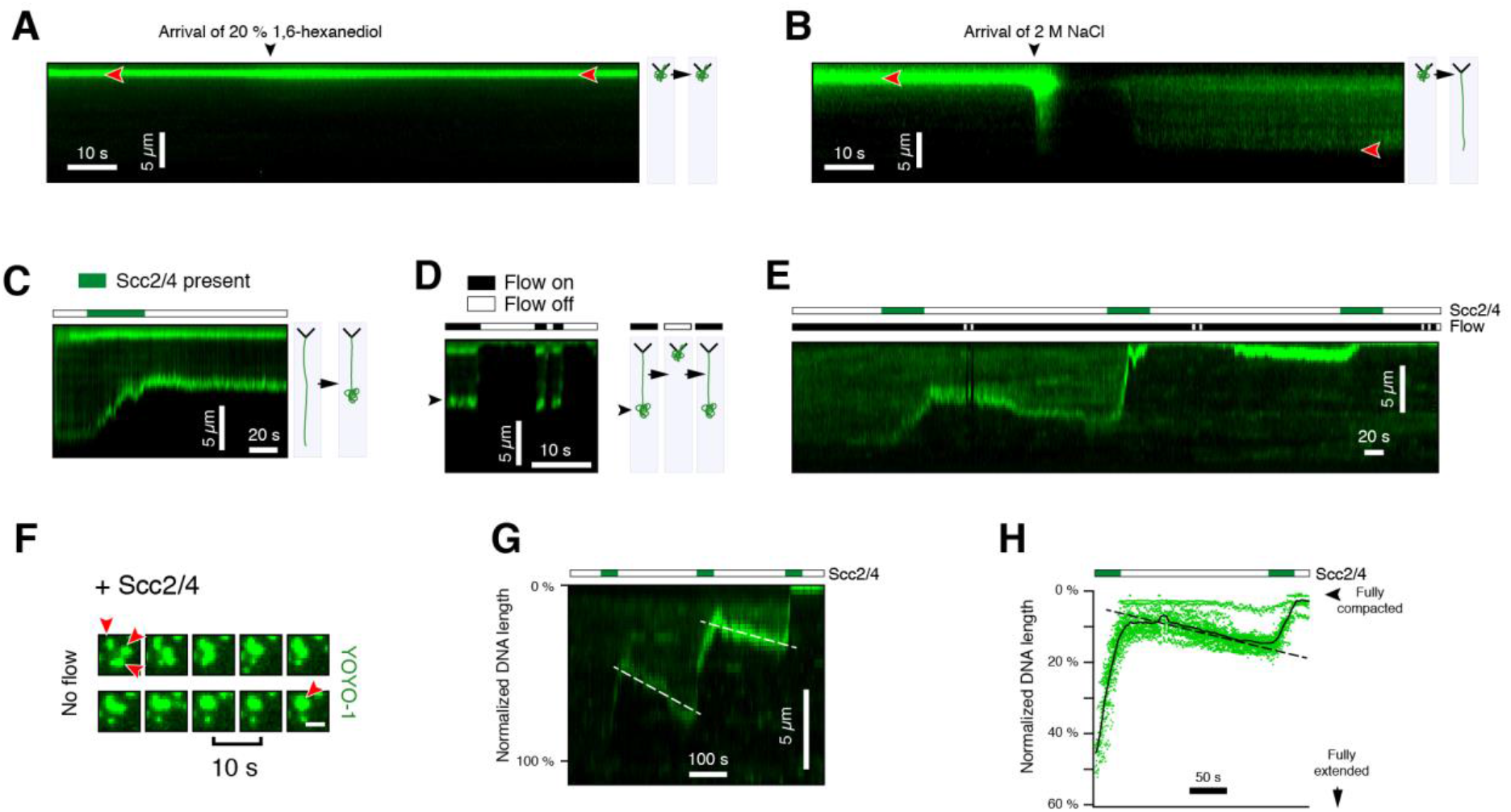
DNA condensation and decondensation dynamics by Scc2/4. (**A+B**) Kymograms of an Scc2/4-induced condensate on a single DNA molecule, challenged with 20% 1,6-hexanediol (**A**) or 2 M NaCl (**B**). Red arrows indicate the free end. (**C**) Kymogram of flow-stretched DNA upon injection of a short pulse of 100 nM Scc2/4. The DNA condenses only while protein is in the flow chamber (green bar indicates protein presence). (**D**) In the absence of Scc2/4 in the flow chamber, DNA does not condense further when it is allowed to relax by relieving the flow tension. (**E**) Kymogram of a multi-pulse experiment. The DNA only condenses while Scc2/4 is in the buffer chamber (indicated by green bar). (**F**) Multiple mobile lipid-anchored DNA molecules coagulate to form larger condensates while Scc2/4 is present. (**G**) Condensation clusters are reversible. DNA compacts while Scc2/4 is present in the buffer chamber and slowly decompacts in the absence of Scc2/4. White lines indicate the end of the DNA molecule. (**H**) Decondensation of 52 DNA molecules between pulses of Scc2/4. Black line: mean. Dashed line: decondensation trend to guide the eye.

Contrasting the LLPS model, our findings are instead compatible with the electrostatics-dependent mechanism of polymer-polymer phase separation (PPPS, (Erdel & Rippe, 2018)), which entails the bridging of polymers (DNA) through multivalent proteins (Scc2/4). As the PPPS mechanism additionally predicts that free protein is required to induce the formation of condensates we tested this possibility in pulsed injection experiments, where we injected short 50 μl pulses of 30 nM Scc2/4 onto a dsDNA curtain that was stretched at a flow rate of 0.1 ml/min. Intriguingly, we found that the condensation of DNA only proceeded while Scc2/4 was present in the flow cell and stopped after free Scc2/4 had been washed out (**Fig. 3C**), resulting in a DNA-Scc2/4 co-condensate at the end of the DNA. We tested if condensates at the end of the DNA are capable of recruiting additional DNA, even in the absence of free Scc2/4, by stopping the flow and allowing the DNA to relax. However, we found that in repeated flow on/off cycles, the length of the condensed DNA remained the same (**Fig. 3D**). While condensate sizes remained stable in absence of free Scc2/4, condensation readily restarted when additional pulses of free Scc2/4 were injected into the flow cell (**Fig. 3E**) and existing condensates were then also able to fuse (**Fig. S2D**). We also observed merging and coalescence of freely diffusing lipid-bilayer-anchored DNA-Scc2/4 condensates in presence of free Scc2/4 (**Fig. 3F**). Hence, we conclude that the formation of Scc2/4-mediated DNA condensates requires free Scc2/4 as a DNA bridging factor, which fulfills the characteristics of PPPS.

### Condensates are dynamic and reversible

A biologically relevant Scc2/4-mediated DNA and chromatin condensation mechanism must be reversible. In the previously described experiments we have shown that condensates are static in the absence of protein, i.e., they do not condense further. When we exposed partially condensed DNA to long washes with buffer in the absence of free Scc2/4, we observed reproducible slow decondensation and concurrent lengthening of DNA (**Fig. 3G**, white dashed line). DNA condensed in the presence of Scc2/4 and decondensed in its absence, indicating that Scc2/4-mediated DNA condensates are thermodynamically reversible. In a buffer flow of 0.1 ml/min, the decondensation rate was ∼3%/min (**Fig. 3H**).

### Condensation is primarily driven by random collisions

DNA compaction as a precursor to condensate formation can occur through loop formation by active, energy-consuming translocation of DNA binding factors (Goloborodko et al., 2016), but also by passive mechanisms like DNA bridging, bending or wrapping (Johnson et al., 2008). The cohesin loader Scc2/4 has no known nucleotide hydrolysis activity and we found no significant influence of ATP on DNA compaction by Scc2/4 (**Fig. S2C**), ruling out active processes. We, therefore, further explored the possible mechanisms for passive DNA compaction. As in previous experiments, we used DNA curtains and exposed single-tethered, flow-stretched 48.5 kbp DNA molecules to near-physiological concentrations of Scc2/4. We followed the DNA compaction over time by observing the accumulation of the YOYO-1 signal. A kymogram of a typical compaction curve for 100 nM Scc2/4 (**Fig. 4A**) shows the appearance of a bright spot at the free end of the DNA, which gradually moves closer towards the tethered end. This indicates that the compaction primarily starts at the free end. Intensity profiles along the DNA at various time points (**Fig. 4B**) also confirm the appearance of a DNA condensate at the free end. Interestingly, when we measured the fluorescence intensity upstream of the compacting end (**Fig. 4C**), we found that it remained not constant but increased almost linearly as the DNA compacted (**Fig. 4D**). This indicates that there is also internal compaction distant from the free end.

**Figure 4.**
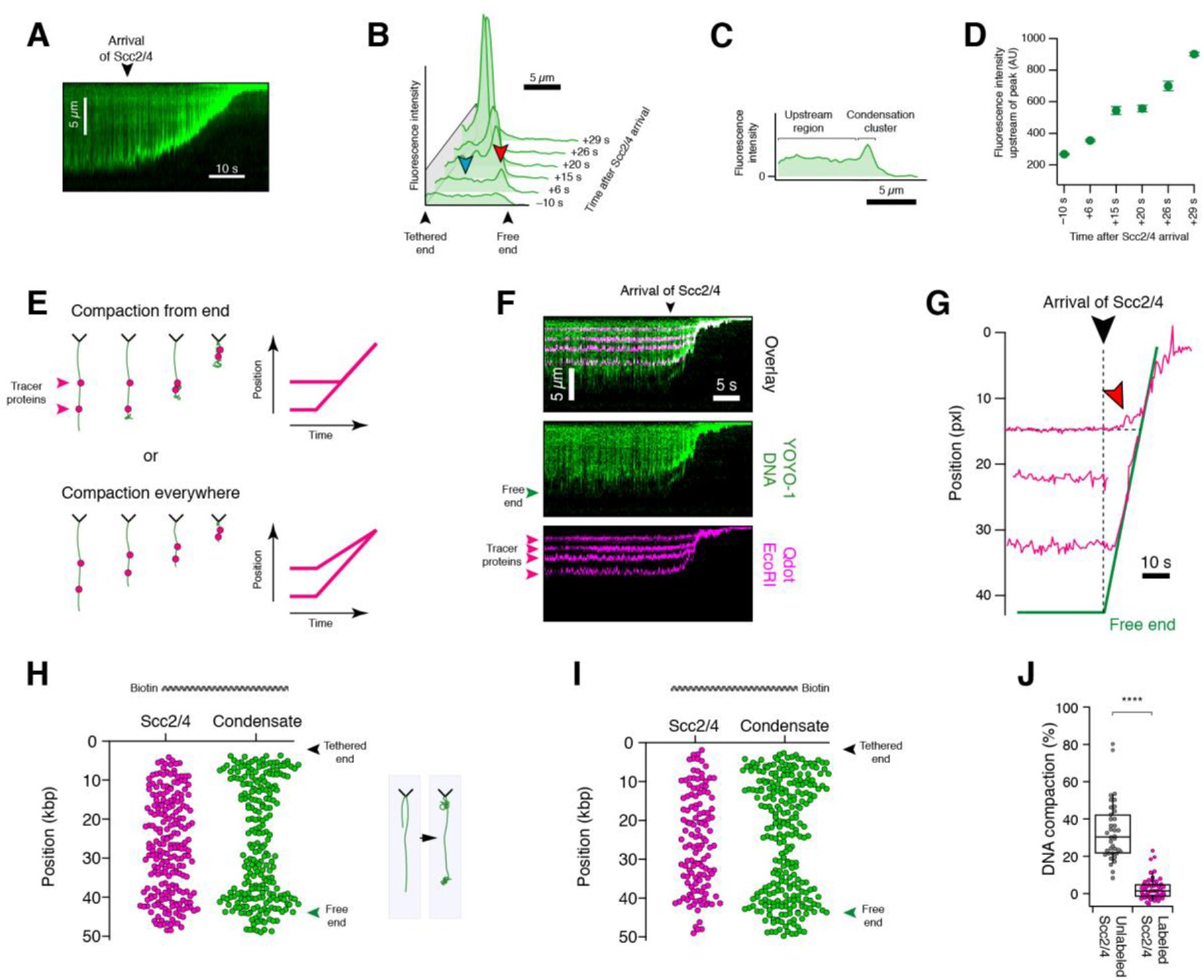
Direction of DNA compaction. (**A**) Kymogram of Scc2/4-mediated DNA condensation shows the appearance of a bright spot at the free end. (**B**) YOYO-1 intensity profiles of a single lambda DNA molecule during Scc2/4-mediated condensation. Time marks indicate the time after the start of Scc2/4 arrival in the flow cell. Red arrow: Condensate at the free end. Blue arrow: Region upstream of condensate. (**C**) YOYO-1 intensity profile along a single lambda DNA molecule during Scc2/4-mediated condensation. (**D**) Average YOYO-1 intensity of the region upstream of the free end (see blue arrow in (**B**)). (**E**) Models for the condensation of DNA. (**F**) Kymograms of Scc2/4-mediated condensation of lambda DNA, which was marked with fluorescently tagged EcoRI-E111Q tracer proteins. (**G**) Tracked positions of tracer proteins upon Scc2/4 addition. The red arrow shows movement of an internal tracer far away from the condensing end. (**H**) Binding positions of Qdot-tagged Scc2/4 (magenta) show a homogenous binding preference across the substrate. Condensation clusters (green) primarily occur at the free end as well as close to the barriers. Data points were pooled from two independent experiments. (**I**) Experiment as in (**G**), with DNA that was tethered at the opposite end. Data points were pooled from three independent experiments. (**J**) DNA compaction of 30 nM unlabeled or labeled Scc2/4. For unlabeled and labeled protein binding, 38 and 50 DNA strands were analyzed, respectively. Distribution is according to student’s t-test significantly different.

To more thoroughly distinguish potential mechanisms of compaction from the end (e.g., through the capture of randomly fluctuating loops) or compaction everywhere (e.g. by bending), we used tracer proteins to track the position of various positions along the DNA during compaction (H. Kim & Loparo, 2016). Generally, in a scenario where compaction occurs solely from the end, we expect that the inner tracer molecules are static until they encounter the compacting end (**Fig. 4E**). On the contrary, a scenario where compaction occurs everywhere predicts movement of internal tracers as soon as the DNA is exposed to the compacting factor. **Fig. 4F** shows a representative kymogram of the Scc2/4-mediated compaction of lambda DNA with Qdot-labeled EcoRI-E111Q tracer molecules bound. Tracking of the tracer molecules then allowed us to follow the compaction path (**Fig. 4G**). To our surprise, we found that, while our observations primarily corresponded to the scenario of compaction from the end, the internal tracer proteins also started moving as soon as the DNA is exposed to Scc2/4 (red arrow in **Fig. 4G, Fig. S4B**). This indicates that while capture of freely fluctuating DNA loops at the free end is the primary mechanism for DNA compaction and condensation, there is also a smaller contribution of internal compaction, perhaps through DNA bending.

We further considered the possibility that the movement of internal tracer molecules could also be a consequence of altered hydrodynamic properties of the DNA during apparent shortening in flow. To address this question, we turned to coarse-grained molecular dynamics simulations where lambda DNA is represented by a Rouse model of 160 beads that were connected by segments of ∼300 bps in length while internal compaction was excluded. After applying flow stretching forces (**Fig. S4C, left**), we modeled the effect of Scc2/4 by introducing an attractive Lennard-Jones-like potential between the beads (see Methods), which explicitly did not induce a shortening of segments but nevertheless led to a rapid compaction of the modeled DNA (**Fig. S4C, right**) due to condensation at the free end. In contrast to DNA curtain experiments, these simulations did not display the movement of internal tracer molecules until they were encountered by the condensing end. This suggests that the experimentally observed movement of tracers is not caused by a change in hydrodynamics but indeed by internal compaction or bending of DNA initiated by Scc2/4 binding.

Finally, we asked if end-dependent compaction may be caused by preferred association of Scc2/4 to free ends. To this end, we Qdot-labeled Scc2/4 and tracked the binding position of Scc2/4 before the onset of condensation (**Fig. 4H**). While DNA condensates initially formed primarily at the free end, the binding position of Scc2/4 showed no preference. We obtained the same result when we anchored the DNA molecule at the opposite end suggesting sequence-independent binding (**Fig. 4I**). In both orientations we also observed a fraction of condensate clusters close to the tethered end, which we attribute to the presence of shorter DNA molecules. However, while Qdot-labeled Scc2/4 was clearly able to bind DNA, labeling had a partially disruptive effect on DNA-compaction (**Fig 4J**), plausibly through interfering with at least one of Scc2/4’s DNA binding sites.

### Scc2/4 compacts DNA by a tension-dependent bridging mechanism

To unravel what drives Scc2/4-mediated bridging on DNA, we utilized optical tweezers and first studied the influence of DNA tension on Scc2/4 interaction. A single lambda DNA molecule was tethered between two optically trapped beads, which allows force application and measurement in a very accurate manner. We incubated the DNA with 10 nM of Scc2/4 first in a tensioned (7.5 μm bead distance) and then in a more relaxed conformation (5 μm bead distance, **Fig. 5A, Fig. S5B**) and found in both cases a strong gradual increase in force due to protein association to the DNA (**Fig. 5B** magenta trace, see also **Fig. S5A** for responses of individual tethers). Naked DNA (green) did not show a change in force, while DNA that was transferred into buffer after protein incubation (black) displayed a slight decrease in force due to unbinding of protein (**Fig. 5B**). To quantitate and compare the degree of compaction at different tension, we calculated the apparent contour length of the tethers as a function of time. Although compaction occurred at both incubation steps, Scc2/4-mediated compaction was much stronger when DNA was held at 5 μm (**Fig. 5C**), where DNA is more relaxed and can be bridged much easier (**Fig. 5A**).

**Figure 5.**
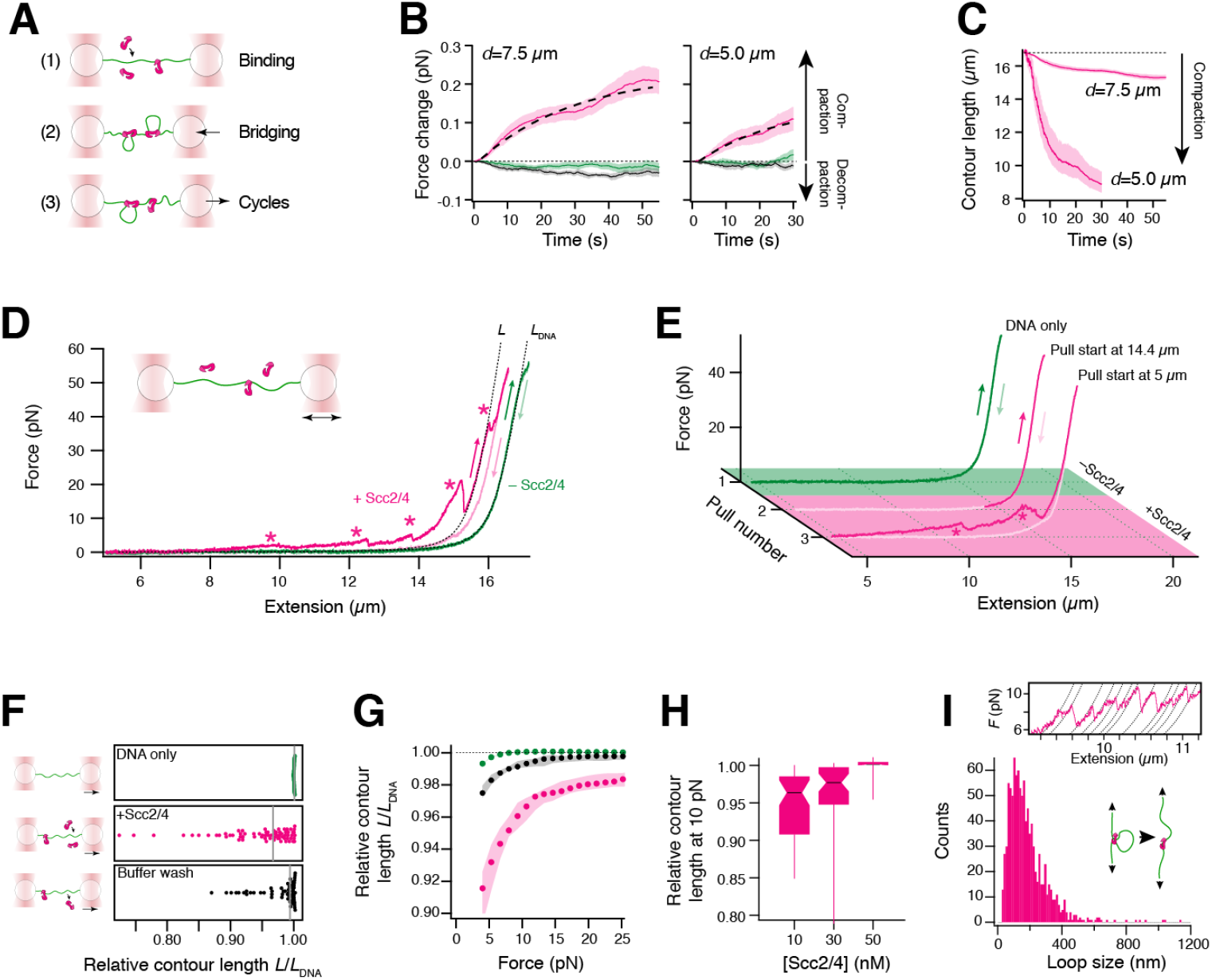
Optical tweezers experiments for understanding the DNA coagulation mechanism of Scc2/4. (**A**) Schematic representation of the experimental procedure. (**B**) Transient force response of DNA tethers held at the indicated distance in flow channels without Scc2/4 (green), with 10 nM Scc2/4 (magenta) or in a buffer channel after previous Scc2/4 incubation (black). Colored line represents the mean of 15 tethers, shading represents the SEM, dashed line represents an exponential fit. (**C**) Time-dependent reduction of the tether contour length in the presence of 10 nM Scc2/4. (**D**) Force-extension curves (FECs) of lambda DNA in the absence (green) and presence (magenta) of 10 nM Scc2/4. Darker shades are stretch traces, lighter shades are relaxation traces. Dashed lines indicate the expected force response of a full-length DNA or a DNA that is shortened by partial compaction, based on an eWLC model. (**E**) Subsequent pulls of lambda DNA that was transferred from the buffer channel (green) to the channel with 10 nM Scc2/4 (magenta). The DNA was incubated in the protein channel in an extended state (pull 2) or at a lower tension (pull 3). Observed compaction is marked by asterisks. (**F**) Relative contour length of DNA stretch-traces from multiple molecules at 10 pN, normalized to the expected contour length of lambda DNA. Green: In the absence of Scc2/4, magenta: In the presence of 10 nM Scc2/4, black: In the buffer channel, after previous Scc2/4 incubation. Grey lines indicate the median. (**G**) Median relative contour lengths of stretch traces at various forces. Shading indicates 68% confidence intervals from bootstrapping. Color code as in (**F**). (**H**) Relative contour lengths at 10 pN after incubation at different protein concentrations. Number of analyzed tethers 10nM: 15, 30 nM: 10, 50 nM: 13. (**I**) Determination of the length of opened loops from sawtooth features. Top: representative stretch FEC with eWLC fits to individual features. Bottom: histogram of loop lengths of 16 individual DNA tethers incubated with 10 nM Scc2/4.

Next, we wondered whether the condensates can be removed from DNA by increasing forces. We applied five stretch and relax cycles to the DNA and recorded force-extension curves (FEC). We observed a Scc2/4-dependent sawtooth pattern, which is a key characteristic of protein-mediated DNA bridges that are ruptured at increasing forces (**Fig. 5D**). While most bridges could be resolved by moderate tensions, we also observed bridges that resisted at forces up to ∼65 pN, where the DNA undergoes force-induced melting. Note, that broken bridges did not reform during relaxation of the DNA strand (**Fig. 5D**), but readily re-formed on fully relaxed DNA, as evidenced by the reappearance of the sawtooth pattern in subsequent cycles (**Fig. S5C**). Additionally, we recognized the absence of the sawtooth pattern, when DNA was incubated at very high tension (14.4 μm bead distance) and directly stretched (**Fig. 5E**). These results indicate that bridging requires relaxed DNA, but does not necessitate the presence of a free DNA end.

We characterized the observed shortening of the DNA by comparing the relative DNA contour length (L/L_DNA_, see **Fig. 5F**) of naked DNA with DNA incubated in Scc2/4 and observed clear DNA compaction in presence of Scc2/4, which was reduced to almost naked DNA level when washing the DNA in buffer after protein incubation (**Fig. 5G**). When looking at individual cycles, we found the strongest compaction for the first cycle, caused by the prior incubation step, while the other four cycles showed reduced but constant compaction as proteins constantly bind and unbind (**Fig. S5D**). During the buffer wash, compaction is reduced in each cycle due to continuous protein unbinding (**Fig. S5D**). We conclude that DNA compaction by Scc2/4 is reversible and can be removed by buffer washes or mechanical tension.

Besides DNA tension, it is likely that also protein concentration influences DNA compaction. To quantitate the dependency on protein concentration, we recorded FECs at increasing Scc2/4 concentrations and found, surprisingly, that compaction was reduced at 30 nM Scc2/4 and completely vanished at 50 nM (**Fig. 5H**). Similarly, we also found that, during incubation, higher protein concentrations counterintuitively resulted in less compaction (**Fig. S5E**). While this finding is puzzling at first glance, it is compatible with the PPPS model of multivalent Scc2/4 bridging DNA: In the case of high protein coverage, there are no more freely accessible DNA binding sites and additional bridges are abrogated, resulting in less DNA compaction.

Finally, to quantify the size of DNA loops formed by bridging, we fitted extensible worm-like chain (eWLC) models to individual rupture events in the sawtooth pattern of FEC and collected all loop sizes in a histogram (**Fig. 5I**). The mean loop size of 183 ± 4 nm (∼540 nt) suggested that DNA is not just simply wrapped around Scc2/4 but bridged over longer distances.

### Scc2/4-DNA co-condensates are preferred loading sites for cohesin

The stark compaction and condensation properties of Scc2/4 raise the possibility that the observed behavior has direct consequences for Scc2/4’s role as a loading factor for cohesin. To address this question, we performed DNA curtain experiments where we pre-loaded 30 nM Scc2/4 to initiate condensate formation followed by loading of 20 nM Qdot-labeled tetrameric cohesin (SMC1, SMC3, SCC1, SCC3) with a continuous flow of 0.1 ml/min (**Fig. 6A**). After washing out all unbound protein, we observed in ∼70% of all cases that cohesin bound directly to Scc2/4-DNA co-condensates while the other proteins attached to uncompacted regions of the DNA (**Fig. 6B**). Kymograms of the whole binding process revealed that, apart from direct binding to the condensate or free DNA, some proteins also initially associated to uncompacted DNA and then slid towards the condensates to stick there (**Fig. 6D**). When incubating cohesin and Scc2/4 simultaneously for 10 min on DNA curtains we observed cohesin overwhelmingly enriched at condensates (**Fig. S6A**). This suggests that most cohesins that initially associated to uncompacted DNA did either dissociate or diffuse towards condensates during incubation. In contract, when applying Qdot-labelled cohesin to a non-compacted, Scc2/4-free DNA curtain (**Fig. S6C**), we observed eight times fewer binding events (**Fig. 6C**). These results show that cohesin is preferably loaded on DNA via association with Scc2/4-formed condensates and that binding is strongly enhanced through this process.

**Figure 6.**
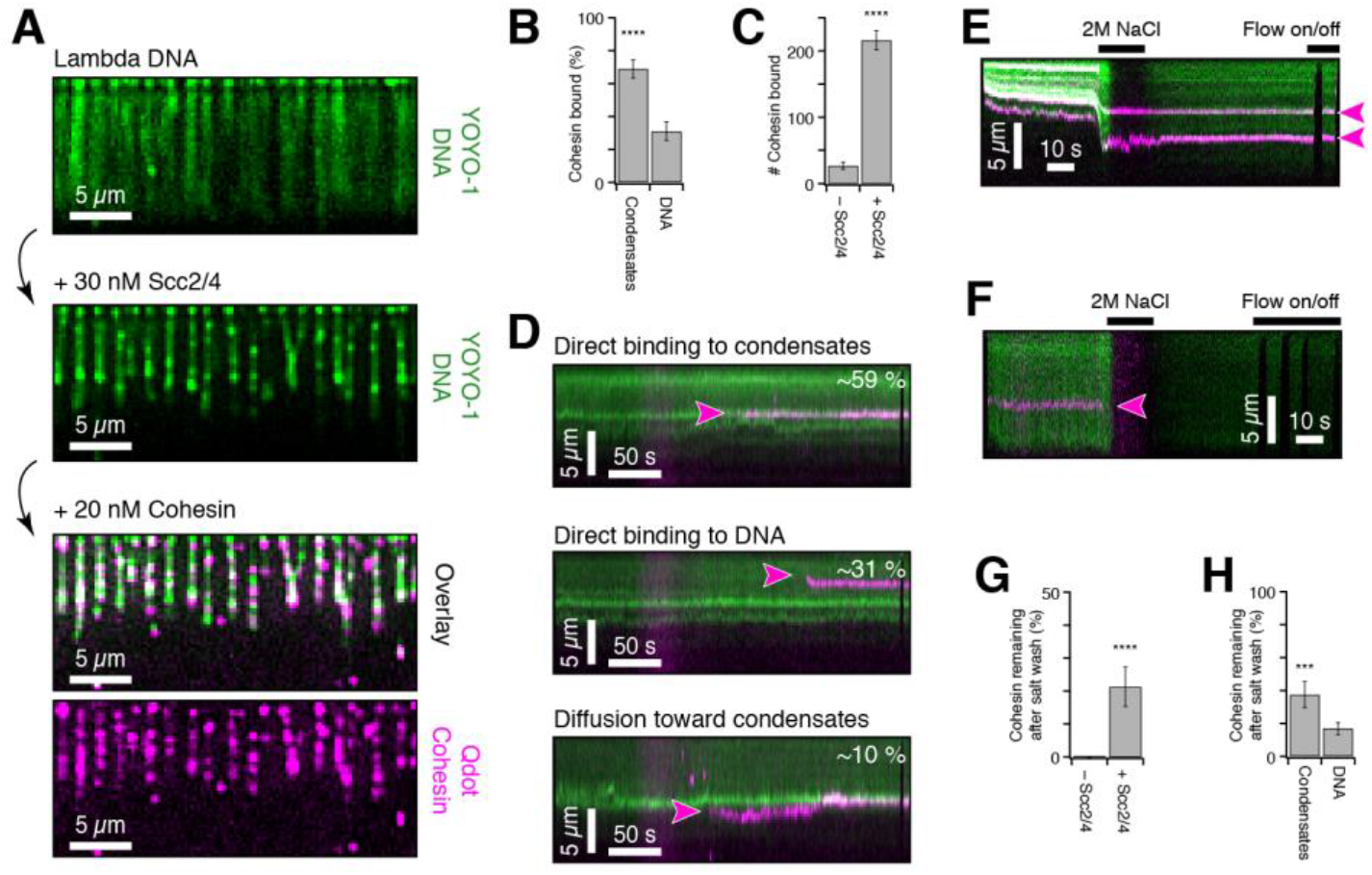
Scc2/4-mediated DNA condensates lead to enhanced loading of cohesin. (**A**) DNA curtain experiment of Qdot-labeled cohesin (magenta) on Scc2/4-mediated DNA condensates (green). (**B**) Ratio of bound cohesin at condensates or at uncompacted DNA after unbound protein was washed out. (**C**) Number of bound cohesin in absence or presence of Scc2/4. (**D**) Representative kymograms of cohesin binding directly to condensates, directly to uncompacted DNA or diffusing towards condensates. (**E**) Kymogram of 2M NaCl wash after cohesin was loaded in presence of Scc2/4. Cohesin remains bound while DNA is decompacted and clusters are dissolved. (**F**) Kymogram of 2M NaCl wash after cohesin was loaded in absence of Scc2/4. Cohesin cannot stay bound on DNA during salt wash. (**G**) Proportion of cohesin molecules remaining on the DNA after 2 M NaCl wash when loaded with or without Scc2/4. (**H**) Proportion of cohesin molecules remaining on the DNA after 2 M NaCl wash when loaded in presence of Scc2/4 at condensates or uncompacted DNA. (**B**+**C**+**G**+**H**) Data was pooled from three independent experiments, error bars represent the standard error of a binominal distribution. Significance was calculated by a z-test.

Furthermore, we tested whether the recruitment observed in this assay constitutes bona fide loading of cohesin. In vitro loading of cohesin had been tested and investigated by performing washes with high concentrations of monovalent salt (Gutierrez-Escribano et al., 2019; Minamino et al., 2018; Murayama & Uhlmann, 2014; Ryu et al., 2021). Recapitulating these experiments, we washed the DNA after loading with 2M NaCl. As expected, we observed decondensation together with stretching of the DNA (**Fig. 6E**). The salt wash removed 79 % of the bound cohesin complexes. However, 21 % resisted the high salt conditions, indicating a strong interaction mechanism (**Fig. 6G**). In absence of Scc2/4, high salt washes removed the bound cohesin complexes completely (**Fig. 6F,G**) showing that Scc2/4 effectively promotes the formation of salt-stable cohesin-DNA interactions. Intriguingly, cohesin that had loaded at Scc2/4-DNA co-condensates exhibited a significantly higher resistance to salt washes (38 ± 8 % remaining) than cohesin that had loaded at uncondensed DNA (17 ± 4 %) (**Fig. 6H**). We conclude that Scc2/4-mediated DNA co-condensates serve as preferred sites for stable cohesin loading.

## Discussion

Here, we have described and characterized a previously unreported property of the cohesin loader Scc2/4. While co-condensation of Scc2/4 with DNA may be unexpected, phase separation phenomena in the context of cellular and nuclear organization are, in fact, common. In the past years it has emerged that phase separation is an important feature to organize the crowded interior of cells, to separate counteracting reactants but also to pool interacting molecules (Erdel & Rippe, 2018). Besides the formation of distinct phase separated bodies, like stress granules, p-bodies or nucleoli (Verdile et al., 2019), phase separation also promotes chromatin organization (Gallego et al., 2020; Larson et al., 2017; Strom et al., 2017) and regulation of processes like transcription (Boehning et al., 2018; Boija et al., 2018; Kent et al., 2020; Sabari et al., 2018; Wei et al., 2020), replication (Parker et al., 2019) or DNA repair (Kilic et al., 2019). In fact, the cohesin complex phase separates in presence of the loader protein Scc2/4 (Ryu et al., 2021). Furthermore, ParB, a bacterial SMC loading factor (Minnen et al., 2011; Sullivan et al., 2009), undergoes phase separation in vivo (Broedersz et al., 2014; Guilhas et al., 2020) and in vitro (Madariaga-Marcos et al., 2019) suggesting that condensate formation might be a general feature of SMC loading proteins.

### Phase separation vs. aggregation

In this work, we have demonstrated that the loading complex Scc2/4 is able to bridge and compact DNA into protein-DNA co-condensates independently of cohesin. Previous studies did not investigate full-length Scc2/4 alone and have used protein concentrations in the low nanomolar range leading to only rarely observed protein clusters, which were often attributed to aggregation or impurities (Davidson et al., 2019; Gutierrez-Escribano et al., 2019; Y. Kim et al., 2019). Our findings extend to physiological protein and salt concentrations (300 nM Scc2/4 (Ho et al., 2018), 150 mM NaCl), strongly suggesting that the observed phenomenon is also relevant in vivo. The presented results are compatible with well-known hallmarks of phase separation (Mitrea et al., 2018), as the condensates exhibit liquid-like behavior represented by droplet formation in bulk (**Fig. 2A**), fusibility in the presence of protein (**Fig. 3F, Fig. S2D**) and decomposition of condensates caused by exchange with the environment (**Fig. 3G**). Condensate dynamics and reversibility also rule out the possibility that condensation may be triggered by protein aggregation.

### Protein constituents for generating phase separation

Many phase separating proteins contain intrinsically disordered regions (IDR) which promote high levels of self-interaction and crowding (André & Spruijt, 2020) and lead to liquid-liquid phase separation (LLPS). Scc2/4 lacks IDRs but comprises HEAT-repeats, which were found before to induce phase separation (Aktar et al., 2019) due to their amphiphilic and very flexible structure (Kappel et al., 2010; Yoshimura & Hirano, 2016). However, we propose that the observed Scc2/4-DNA co-condensates are formed by polymer-polymer phase separation (PPPS), where Scc2/4 serves as a multivalent bridging factor for DNA and does not self-interact. In our droplet experiments, both polymers, DNA and Scc2/4, are required to form liquid-like droplets (**Fig. 2A**), which corroborates a PPPS-based mechanism. PPPS has previously been shown for the related SMC loading factor ParB, which bridges DNA by multiple helix-turn-helix motifs and also does not self-interact (Fisher et al., 2017; T. G. W. Graham et al., 2014). DNA decondensation during salt wash (**Fig. 3B**) revealed that DNA bridging is promoted by electrostatic rather than hydrophobic interactions (Erdel & Rippe, 2018). Furthermore, we found that free protein is required for condensate formation and coagulation (**Fig. 3E**; **Fig. 5G**), as it is driven by the introduction of new bridging factors until the binding sites are saturated (Erdel & Rippe, 2018). Also other DNA compacting proteins require free protein in solution for DNA condensation, which is thought to facilitate dynamic and flexible DNA binding and therefore influence gene regulation (J. S. Graham et al., 2011). In contrast to that, the LLPS forming protein HP1α continuously compacts DNA in absence of free protein, only triggered by pre-bound HP1α (Keenen et al., 2020). DNA compaction frequently occurs in DNA curtain experiments at Scc2/4 concentrations higher than 10 nM (**Fig. 2C**), which suggests that the DNA needs to be coated with protein to a certain extent to induce condensate formation through a coil-globule transition (Lappala & Terentjev, 2013; Quail et al., 2020; Shendruk et al., 2015). However, our optical tweezer data show no linear relation between protein amount and compaction behavior as higher concentrations prevent compaction dramatically (**Fig. 5H**). This might be explained by a reduction of possible DNA bridging points when DNA is completely coated by Scc2/4 at concentrations higher than 10 nM and confirms the existence of at least two independent DNA binding sites in Scc2/4 (Kurokawa & Murayama, 2020). HU protein was shown before to exhibit a comparable behavior compacting DNA at low concentration by bending and reextending DNA at higher concentrations due to protein filament formation (Song et al., 2016). The discrepancy between critical protein concentrations for DNA condensation in DNA curtain and optical tweezer experiments might be due to different incubation procedures as well as different effective DNA to protein ratios in the assays.

### Bridging vs. bending

Besides protein-mediated DNA bridging, DNA compaction can also be caused by DNA bending which alters DNA flexibility and regulates replication, recombination and gene expression (Dillon & Dorman, 2010; Gruber, 2014). Our optical tweezer experiments clearly show a sawtooth pattern in FECs (**Fig. 5D**), which is a known characteristic of DNA looping (Dame et al., 2006) as every rip during DNA stretching marks a broken protein-DNA bridge. In our DNA curtain experiments, Scc2/4 binds sequence-independently to the lambda DNA (**Fig. 4H,I**) (Taylor et al., 2015), but nevertheless, compaction always starts from the free DNA end (**Fig. 4A,B**). This was also reported for other DNA binding proteins, like HP1α (Keenen et al., 2020), FUS (Renger et al., 2021) or the transcription factor Klf4 (Morin et al., 2020) as well as for ParB (T. G. W. Graham et al., 2014) and can be explained by a preference of these proteins to bridge where tension is lowest and DNA is highly flexible (Baumann et al., 2000; H. Kim & Loparo, 2016). Optical tweezer experiments confirmed this by increased compaction at lower tension (**Fig. 5C,E,G**) as well as the fact that condensates do not re-form during DNA relaxation but only at minimal tension at the beginning of each cycle (**Fig. 5D, Fig. S5C**). All these findings suggest a clear compaction mechanism driven by bridging. However, our fluorescence intensity (**Fig. 4C**) and tracer protein studies (**Fig. 4G, Fig. S4C**) suggest also an internal compaction mechanism most likely caused by DNA bending (Carrivain et al., 2012) probably at the helix-turn-helix motifs (Harrington & Winicov, 1994). The contribution of both, bridging and bending, to the compaction mechanism was previously discovered for a bacterial SMC protein (H. Kim & Loparo, 2016). Furthermore, crystal structures as well as molecular models of cohesin in complex with Scc2/4 suggest that DNA bending might be an essential step for cohesin loading (Higashi et al., 2020; Shi et al., 2020). However, in optical tweezer experiments, we could not resolve a reduction in DNA persistence length, which was found to occur during DNA bending (Farge et al., 2012), as the bridging process dominated FEC appearance.

### Loading sites for cohesin

Scc2/4-supported loading of cohesin is more efficient (Stigler et al., 2016) and more stable (Gutierrez-Escribano et al., 2019) than self-loading due to a topological entrapment of the DNA (Murayama & Uhlmann, 2014). Hence, we propose that phase separation might trigger this loading, as we and others could not find condensate formation by cohesin in absence of Scc2/4 (**Fig. S6C**, (Ryu et al., 2021)). Our DNA curtain experiments clearly show that cohesin is recruited preferably to the condensates (**Fig. 6A,B, Fig. S6A**) and that these cohesin molecules are also more resistant to high salt washes (**Fig. 6H**), which suggests a topological entrapment of the DNA (Kurokawa & Murayama, 2020) in a manner that prevents sliding-off the single tethered DNA strand. Interestingly, SMC complexes that load themselves on DNA like MukB (Kumar et al., 2017) and yeast condensin (Yoshimura et al., 2002) also form co-condensates with DNA. Hence, our observations strongly suggest that phase separation induces loading. Protein recruitment to phase separated droplets has also been observed in other contexts. For example, FET-fusion proteins form condensates with DNA and recruit and activate RNA polymerase even without the respective promoter (Zuo et al., 2021). Similarly, the DNA repair protein ATR is recruited to and activated in condensates (Frattini et al., 2021). We therefore propose that also Scc2/4 condensates do not only bind but also activate cohesin, most likely by stimulating ATPase function (Murayama & Uhlmann, 2014; Petela et al., 2018).

### Ideas and Speculation

There are indications that TAD formation and phase separation are counteracting processes (Y. Kim & Yu, 2020), raising the question whether the loading of cohesin at Scc2/4-mediated DNA-condensates determines whether cohesin promotes loop extrusion or sister chromatid cohesion. As cohesin degradation in vivo leads to the loss of all DNA loops but not of the condensation-mediated compartmentalization of chromatin (Rao et al., 2017), it is likely that phase separation does not cause loop extrusion but both processes co-exist. Furthermore, TAD formation clearly depends on ATP and is hence an active process (Gutierrez-Escribano et al., 2019; Y. Kim et al., 2019), while phase separation can organize DNA without energy consumption (Ryu et al., 2021). Sister chromatid cohesion, in contrast, is known to start at replication forks, a very crowded environment, where most likely multiple cohesins are required to resist strong tension (Skibbens, 2000; Zhang et al., 2008). It is possible that loading at condensates sets a starting point to accumulate cohesin and recruit other factors that are required for cohesion maintenance (**Figure 7**). An interesting candidate for such a factor is the cohesin acetylase Eco1, which was previously shown to be recruitable by DNA-protein-co-condensates in other contexts (Zuilkoski & Skibbens, 2020). It is possbile that Scc2/4 is exchanged by these factors within the condensates once its task is fulfilled as it is known that Scc2/4 is not required for cohesion maintenance (Srinivasan et al., 2019). Future studies need to reveal if and how other downstream protein factors are recruited to the condensates and how interchangeable their composition is. Nevertheless, our results provide new insights into the properties and the formation mechanisms of protein-DNA co-condensates and prove the power of single-molecule methods to uncover unexpected features of already well-known protein complexes.

**Figure 7.**
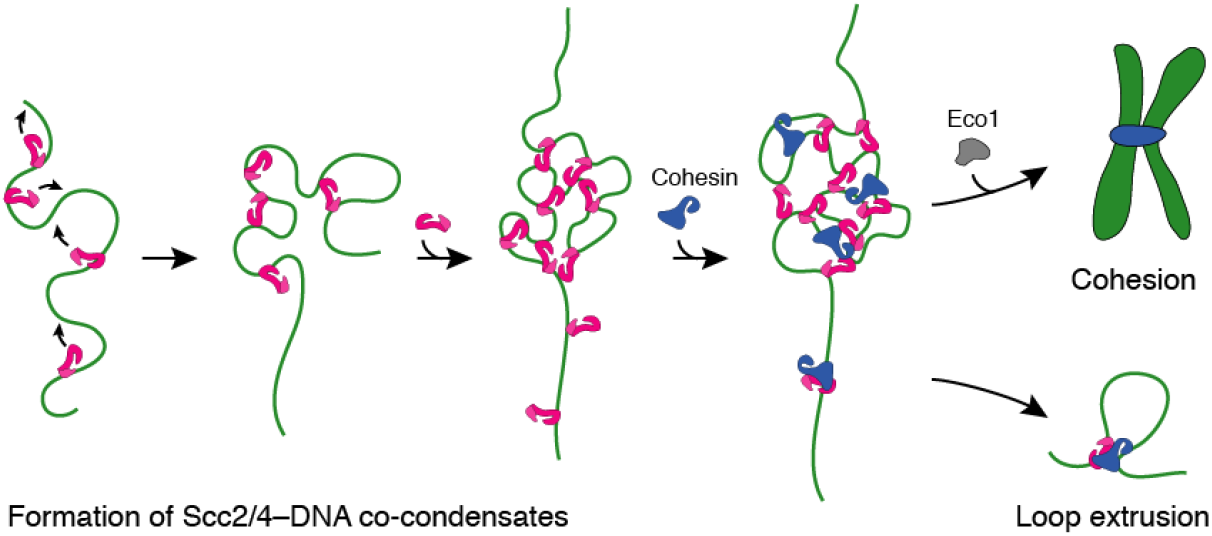
Schematic representation of DNA compaction mechanism. Scc2/4 binds randomly and bridges DNA dependent on DNA tension. Free protein promotes the condensation process by providing new interaction points. Cohesin is preferably recruited to the condensates creating a crowded environment which facilitates the recruitment of other proteins like Eco1 and finally enables sister chromatid cohesion. Cohesin that is loaded on uncompacted DNA might induce loop extrusion.

## Methods

### Protein Expression and Purification

The S. cerevisiae cohesin tetramer (SMC3-SMC1-SCC1-SCC3) and loader complex Scc2/4 were cloned in multicopy episomal vectors (pRS424-GAL1-10-SMC1-Strep-SM3-GAL7-SCC1-HIS-HA + pRS426-GAL-Scc3; pRS426-URA3-GAL-SCC2-3xMyc-3xStrepII-SCC4) and transformed into yeast W303-1a strains. Protein expression and purification was done as described before (Gutierrez-Escribano et al., 2019). In brief, yeast was cultivated at 30 °C in selected dropout medium supplemented with 2 % raffinose and 0.05 % glucose to an OD > 0.8, induced by addition of 2 % galactose and grown for further 20 h at 20 °C. Cells were harvested by centrifugation (4500 rpm, 15 min, 4 °C), washed with PBS and resuspended in buffer A (25 mM Hepes, pH 7.5, 5 % glycerol, 5 mM β-Mercaptoethanol) containing 200 mM NaCl (A-200) and protease inhibitor (cOmplete, Sigma-Aldrich, St. Louis, MI). The cell suspension was frozen dropwise in liquid nitrogen and stored at -80 °C overnight. The frozen yeast suspension was lysed by freezer mill (SPEX sample prep, Metuchen, NY) and afterwards thawed on ice. The solution was supplemented with Pierce Universal Nuclease (0.1 μl/ml, Thermo Fisher Scientific, Waltham, MA) and incubated for 1 h at 4 °C while rotating. After centrifugation (17,000 xg, 30 min, 4 °C) and filtering (0.22 μm, Thermo Fisher Scientific), lysate was purified on a 1 ml Strep Trap HP (GE Healthcare, Chicago, IL) using the Äkta pure purification system (Cytiva, Marlborough, MA) and eluted by buffer A-200 containing 5 mM D-desthiobiotin (Thermo Fisher Scientific). Pooled eluate fractions were adjusted to 150 mM NaCl and purified on a 5 ml HiTrap Heparin column (GE Healthcare) using a linear gradient from 150 mM NaCl to 1 M NaCl in buffer A. Pooled eluate fractions were concentrated, adjusted to 300 mM NaCl and purified on a Superose 6 Increase 10/300 GL (GE Healthcare) using buffer A containing 300 mM NaCl. Eluate was concentrated by 100 kDa Amicon Ultra, frozen in liquid nitrogen and stored at -80 °C.

EcoRI-E111Q mutant was cloned into a pTXB3 vector including an N-terminal 3xFLAG tag and a C-terminal Intein-CBD purification tag (pTXB3-3xFLAG-EcoRI_E111Q-Intein-CBD). For expression, the plasmid was transformed into E. coli Rosetta (DE3) competent cells. Bacteria were cultivated at 37 °C in LB medium until OD_600_ of 0.6 was reached, followed by induction of expression by addition of 1 mM IPTG and further shaking at 20 °C for 4 h. Cells were harvested by centrifugation at 4000 rpm for 10 min, resuspended in 20 ml resuspension buffer (20 mM Tris-HCl pH 7.5, 0.1 mM PMSF, 10 % glycerol) and lysed by sonication. After centrifugation, the supernatant was transferred onto 3 ml of pre-equilibrated (washing buffer: 20 mM Tris-HCl pH 8.8, 0.5 M NaCl) chitin resin, incubated for 30 min at 4 °C and released by gravity flow. The resin was washed with washing buffer and the protein was cleaved overnight at 4°C with 10 ml cleavage buffer (20 mM Tris-HCl pH 8.8, 0.5 M NaCl, 50 mM DTT). The next day, eluate was released and resin was washed with 10 ml washing buffer. Fractions containing EcoRI-E111Q were pooled, dialyzed at 4 °C overnight against storage buffer (40 mM Tris-HCl pH 7.5, 300 mM NaCl, 0.1 mM EDTA, 50 % glycerol, 10 mM β-Mercaptoethanol, 0.15 % Tween20) and stored at -80°C.

### Protein labeling

Scc2/4 was labeled by mixing the protein in an equimolar ratio with Qdot 705 Strep Conjugate (Thermo Fisher Scientific) and incubating for 10 min on ice in BSA buffer (40 mM Tris, pH 7.5, 1 mM MgCl_2_, 1 mg/ml BSA, 1 mM DTT) with 50 mM NaCl. EcoRI-E111Q and cohesin tetramer were labeled with a similar protocol using the anti-Flag Qdot 705 for EcoRI-E111Q and the anti-HA Qdot 705 for cohesin, both produced with the SiteClick Qdot 705 antibody labelling kit (Thermo Fisher Scientific) utilizing monoclonal anti-Flag M2 antibody from mouse (Sigma-Aldrich) and anti-HA high affinity from rat (Sigma-Aldrich).

### Droplet Microscopy

Scc2/4 droplets were imaged on custom-made flow cells using a prism-type total internal reflection microscope (Nikon Ti2e; Nikon, Minato, Japan) with a 60x objective (CFI P-Apochromat VC 60 x WI, Nikon) in brightfield mode. For flow cell production, microscopy slides (borosilicate glass, 76×26 mm, Roth, Karlsruhe, Germany) were equipped with two stripes of parafilm forming a flow channel and covered with a cover slip (18×18 mm, Roth). Parafilm was melted on a 100 °C hot plate to fix the assembly leading to a channel of approximately 15 μl volume. Flow cells were pre-incubated with 40 μl sample buffer (40 mM Tris (pH 7.5), 50 mM NaCl, 1 mM MgCl_2_, 1 mg/ml BSA, 1 mM DTT). Variable concentrations of Scc2/4 were mixed with 30 nM linearized pUC19 (digested with NdeI restriction enzyme and purified by ethanol precipitation) and 0.5 mM ATP in sample buffer (20 μl reaction volume). Protein-DNA mixture was applied to the flow cell and incubated for 5 min. After this, microscopy videos of appr. 20 frames for each sample were recorded in NIS Elements (Nikon) using an electron multiplying charge-coupled camera (Andor iXon life, Andor Technology, Belfast, UK) with illumination times of 100 ms and a sample rate of 200 ms. The background signal (buffer only sample) was subtracted from each video using custom-written software in Igor Pro (WaveMetrics, Lake Oswego, OR). Experiments were performed at least twice for each condition.

### DNA curtain assay

DNA curtains experiments were performed in custom made flow cells produced as described previously (Greene et al., 2010). In brief, Cr barriers were deposited on fused-silica slides by e-beam lithography. The sample chamber around the barriers was confined by double sided tape and covered with a cover slip. The flow cell was connected to a microfluidic system containing a syringe pump and two injection valves by nanoports glued on drilled holes in the fused-silica slide (UQG Optics, Cambridge, UK). Sample was illuminated with a 488 nm laser (OBIS LX/LS, coherent, Santa Clara, CA) in a prism-type total internal reflection microscope (Nikon Ti2e) equipped with an electron multiplying charge-coupled camera (Andor iXon life) with illumination times of 100 ms and a sample rate of 200 ms. Videos were recorded in NIS Elements (Nikon). For single-tethered DNA curtains, 3’-biotinylated λ-DNA (NEB, Ipswich, MA) was attached to the pre-added lipid bilayer surface (DOPC, DOPE, DOPE-biotin, Otto Nordwald, Hamburg, Germany) by biotin-streptavidin-biotin interaction and flow stretched over the Cr barriers to arrange DNA strands in an array. For flipped DNA experiments, 5’-biotinylated λ-DNA was used. For double tethered DNA, double modified λ-DNA (3’- biotinylation and 5’- digoxigenation) was used. After flow stretching of the DNA over the Cr barriers, DNA was anchored to downstream Cr pedestals by digoxigenin binding protein DIG10.3. All experiments were performed in BSA buffer supplemented with 0.16 nM YOYO-1. Different concentrations of Scc2/4 were diluted in BSA buffer containing variable amounts of NaCl (0 mM, 50 mM, 100 mM, 150 mM), 0.5 mM ATP, 100 μM biotin and loaded on the DNA curtain using a continuous flow of 0.1 ml/min for 4 min. For all injections, a 50 μl injection loop and 65 μl sample volume were used. On double tethered DNA curtains, protein was loaded with a flow of 0.3 ml/min and incubated on the DNA for 5 min without flow. After this, unbound protein was flushed out with 0.3 ml/min and remaining protein was documented. The setup for the study of the condensation of mobile surface-tethered DNA was identical to the setup used for DNA curtains, but videos were recorded away from the chromium barriers.

As DNA curtains provide information about multiple DNA molecules at once, sufficient statistics are already achieved with one measurement. We repeated experiments at least once to rule out experimental fluctuations and pooled results from different experiments to one data set. For very high protein concentrations (300 nM) we performed DNA curtains assays only once, as then unspecific protein binding increased massively preventing any further experiments in the flow cell.

For compaction assays with a tracer molecule, 10 nM Qdot-labeled EcoRI-E111Q was supplemented with 0.5 mM ATP and 100 μM biotin in BSA buffer with 50 mM NaCl and loaded on the DNA curtains with 0.1 ml/min continuous flow. After flushing out all unbound protein, 50 nM unlabeled Scc2/4 was flushed into the flow cell and compaction was documented. The experiments were performed in triplicates.

For cohesin recruitment experiments, 30 nM unlabeled Scc2/4 was loaded on the curtains first (in BSA buffer with 50 mM NaCl, 0.5 mM ATP, 100 uM biotin), followed by loading of 20 nM Qdot-labeled cohesin tetramer in the same buffer. For salt washes, 2M NaCl in BSA buffer was applied with a flow of 0.3 ml/min. The experiments were performed in triplicates.

For decondensation, 100 nM Scc2/4 was loaded without laser excitation to prevent photo-crosslinking. After this, 20 % 1,6-hexanediol (Sigma Aldrich) or 2 M NaCl were loaded on the DNA curtains under continuous flow with laser excitation and decompaction was documented. Protein injection profiles were determined by loading blue dextran in BSA buffer on the flow cell and detecting the absorption change over time in brightfield microscopy.

### Data analysis DNA curtains

Videos were analyzed using custom-written software in Igor Pro 8 (WaveMetrics). For determination of compaction (%), DNA length before and after protein exposure was determined manually from kymograms. Spot fluorescence over time was determined with the help of a custom-written multi particle tracker. In cohesin recruitment assays, cohesin binding events as well as the amount of cohesin molecules remaining at the DNA after salt wash were counted in dependence of Scc2/4 presence. Cohesin binding events were also counted according to the binding at loader-formed condensates or at uncompacted DNA. Cohesin stability after salt wash was determined for these subspecies separately. Data was collected in at least two independent experiments and pooled. Standard error was calculated from binomial distribution and significance was determined by a z-test. P-values were calculated from z-values assuming a two-tailed hypothesis and a significance level of 0.05 using Social Science Statistics Web Calculator.

### DNA compaction simulations

The DNA was approximated by a Rouse model of 160 beads, connected by Hookean springs with a length of 100 nm and a spring constant of 0.2 pN/nm. Simulations were run by integrating the time-discretized Langevin equation of the bead motion with a diffusion coefficient of 1.1e^9^ nm^2^/s and a time step of 1e^-9^ s for >=1e^8^ steps. One bead was fixed in space while the other beads experienced a stretching force to extend the DNA to a mean end-to-end distance of ∼10 μm, close to experimental conditions. Condensation conditions that explicitly do not change the equilibrium spacing between the beads were enabled by an additional Lennard-Jones attraction between beads with an equilibrium distance of 100 nm and a potential depth of 5 k_B_T. The position of every 20th bead was recorded. Simulations were run in triplicate.

### Optical tweezers

Optical tweezer experiments were performed on a dual-trap C-Trap (Lumicks, Amsterdam, The Netherlands) including a μFlow microfluidics system and a 4-channel glass slide (Type C2, Lumicks). The system was controlled using the Bluelake software (Lumicks) and custom Python scripts. The first channel of the slide contained 4.36 μm Streptavidin coated polystyrene microspheres (Spherotech, Fulda, Germany) diluted in sample buffer (40 mM Tris, pH 7.5, 1 mM MgCl_2_, 1 mg/ml BSA, 1 mM DTT, 50 mM NaCl). Here, two microspheres were trapped and the stiffness of the optical traps was calibrated using Bluelake’s built-in thermal calibration method. The microspheres were then moved to the second channel which contained 3’,5’-double-biotinylated λ-DNA in BSA buffer while applying a weak flow. The mobile optical trap was moved periodically between a bead distance of 5 and 15 μm while the force acting on the trapped microsphere was observed. Upon detection of the characteristic force increase at increasing bead distances caused by a captured DNA strand, the microspheres were moved to the third channel, which contained sample buffer supplemented with 0.5 mM ATP. There, tethering of a single DNA strand was verified by recording a force-extension curve (FEC) and comparing it to an extensible worm-like-chain (eWLC) model for a single λ-DNA molecule (Wang et al., 1997).

During experiment, the DNA was stretched first to a tension of 0.3 pN and held at this trap separation for 60 s, then stretched to a tension of 0.05 pN and held at this position for 30 s. After this, force-extension curves were recorded for five stretch-relax cycles. This procedure was performed first in the buffer channel, then similarly in the protein channel (variable concentrations of Scc2/4 in sample buffer supplemented with 0.5 mM ATP) and after this again in the buffer channel. For 10 nM Scc2/4, 15 DNA tethers were analyzed by stretch-relax cycles and data points were pooled. For 30 nM and 50 nM Scc2/4, 10 and 13 tethers were analyzed, respectively.

### Data analysis optical tweezers

Analysis of the recorded incubation steps and force-extension curves was performed using custom-written code for the IGOR Pro 8 software (WaveMetrics). The effect of incubation of λ-DNA with Scc2/4 was analyzed by comparing the change in the differential force response over time Δ*F*during the different incubation steps. The constant trap separation d is composed of the sum of DNA extension ξ(*F*)and the bead deflection 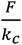,

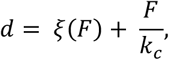

with F = F_t(0)_ + ΔF, and *k*_*c*_ is the trap stiffness. F_t(0)_ is the initial DNA tension, 0.3 or 0.05 pN for the two incubation conditions, respectively. Rearranging the equation yields the DNA extension as

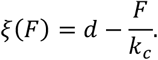

Using an inversion of the eWLC model (Wang et al., 1997), ξ(*F*) can then be converted into a contour length *L*_*c*_ (Mehlich et al., 2020).

For each DNA molecule the force-extension curves (FEC) for naked λ-DNA where fitted with an eWLC model to obtain their persistence length *L*_*p*_ and contour length *L*_*c*_. For all further analysis of the FECs, the persistence length was then fixed to the value obtained for naked DNA. For every stretch FEC, the contour length at each force is then obtained by applying an inverted eWLC to each point of the FEC and binning the result based on the corresponding force value. The contour lengths were then normalized to the previously obtained contour length of naked DNA.

The size of loops formed by Scc2/4 on the DNA was determined using the stretch FECs. Opening of DNA loops during stretching results in a sawtooth pattern. Fitting an eWLC with fixed persistence length to the FEC between two opening events, yields the respective contour length for each sawtooth feature. The loop size is then obtained as the difference in contour length of two consecutive features. Loop sizes were plotted in a histogram, mean was determined from bootstrapping (error = 1-σ).

## Acknowledgments

This work was supported by a DFG Emmy Noether Grant STI673/2-1 and an ERC starting grant (CHROMDOM: 758124). JS acknowledges the support of the LMU Center for Nanoscience CeNS. We thank Isabel Aly for providing EcoRI protein.

## Supplementary Figures

**Figure S1.**
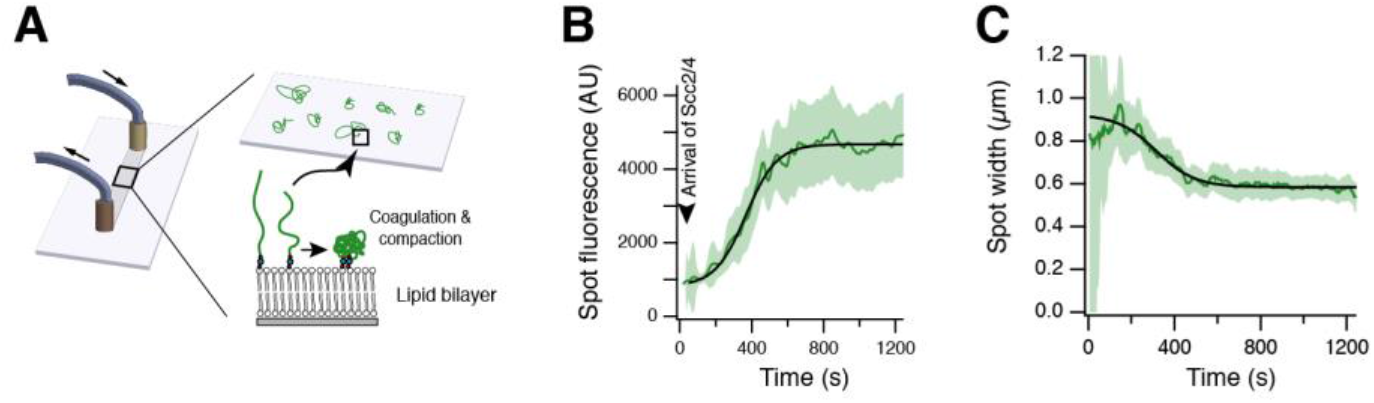
Scc2/4-mediated DNA compaction over time. (**A**) Scheme of flow cell set-up and DNA diffusion on lipid bilayer. (**B**) Fluorescence intensity of freely diffusing DNA spots over time. (**C**) Change in width of freely diffusing DNA spots over time. (**B**+**C**) Dark green curve shows the average of 154 DNA molecules, shading represents 95 % confidence interval.

**Figure S2.**
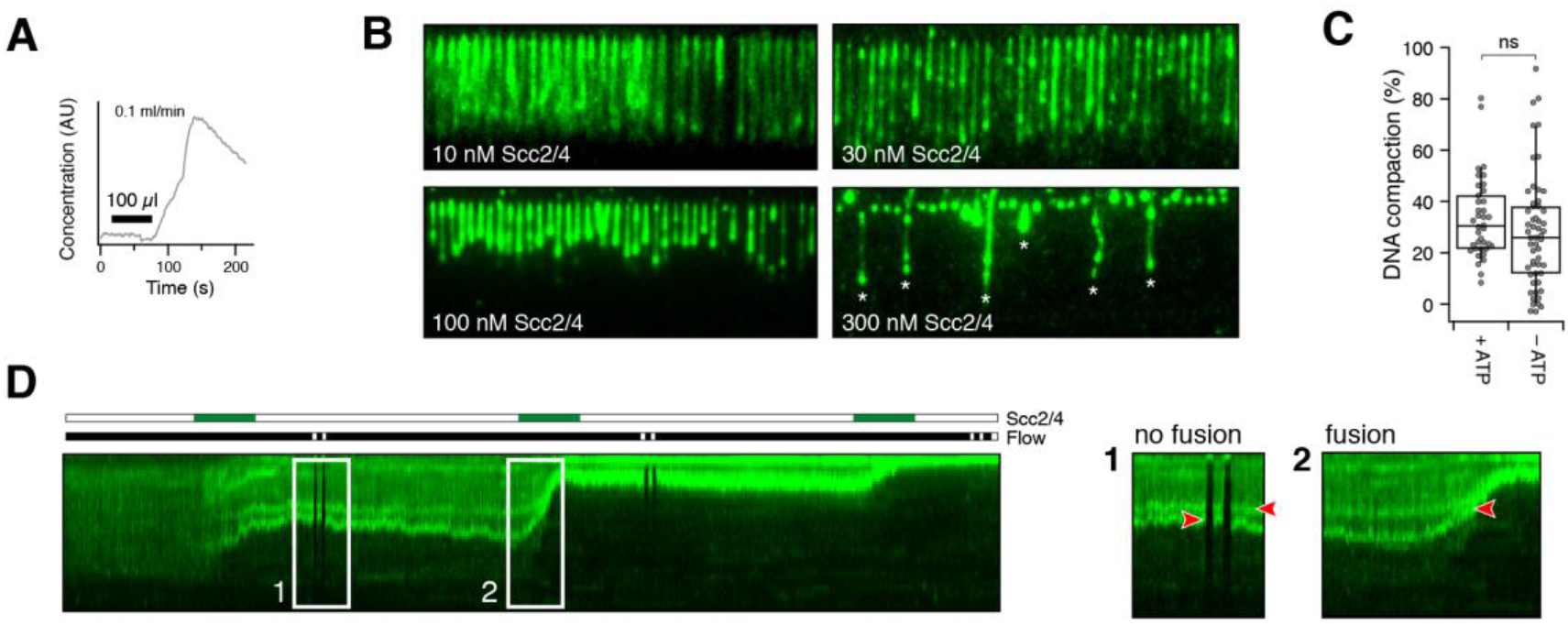
DNA compaction details. (**A**) Injection profile of 50 μl blue dextran with 0.1 ml/min flow measured by change in absorption. Sample arrives after ∼120s in the flow cell, maximum sample concentration: ∼150 s after injection. (**B**) DNA curtain experiments of different concentrations of unlabeled Scc2/4 at 50 mM NaCl to visualize differences in compaction. At 300 nM Scc2/4 some DNA strands are tethered to the lipid surface (asterisks), which were excluded from the analysis. (**C**) DNA compaction mediated by 30 nM Scc2/4 at 50 mM NaCl in presence or absence of 0.5 mM ATP. The means are similar according to student’s t-test (p-value = 0.175), n_(+ATP)_=38, n_(-ATP)_=55. (**D**) Pulsed injection of 3x 50 μl 30 nM Scc2/4. Box 1: Condensates stay separate in absence of Scc2/4 even when the flow is stopped. Box 2: Condensates fuse when free Scc2/4 is present in the flow cell.

**Figure S3.**
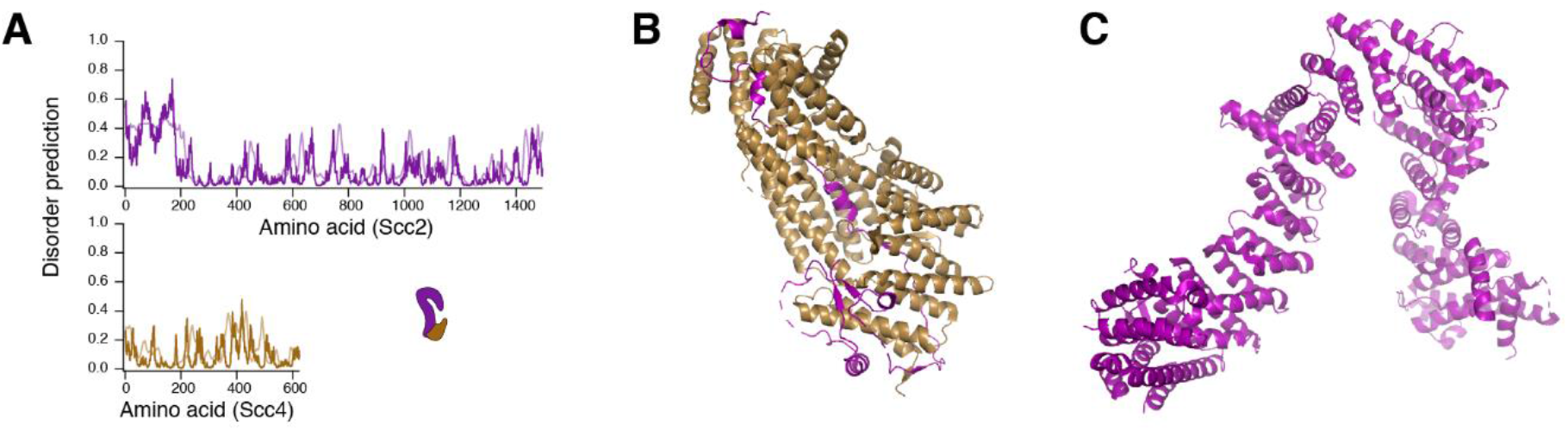
Structural insights into Scc2/4 complex. (**A**) Disorder prediction for Scc2/4 by IUPred2 (dark shading) and ANCHOR2 (light shading) (Mészáros et al., 2018) (**B**) Crystal structure of Scc4 in complex with Scc2(1-181), PDB: 4XDN (Hinshaw et al., 2015). The N-terminal unstructured domain of Scc2 is stabilized by Scc4. (**C**) Crystal structure of Scc2(127-1493), PDB: 5ME3 (Chao et al., 2017).

**Figure S4.**
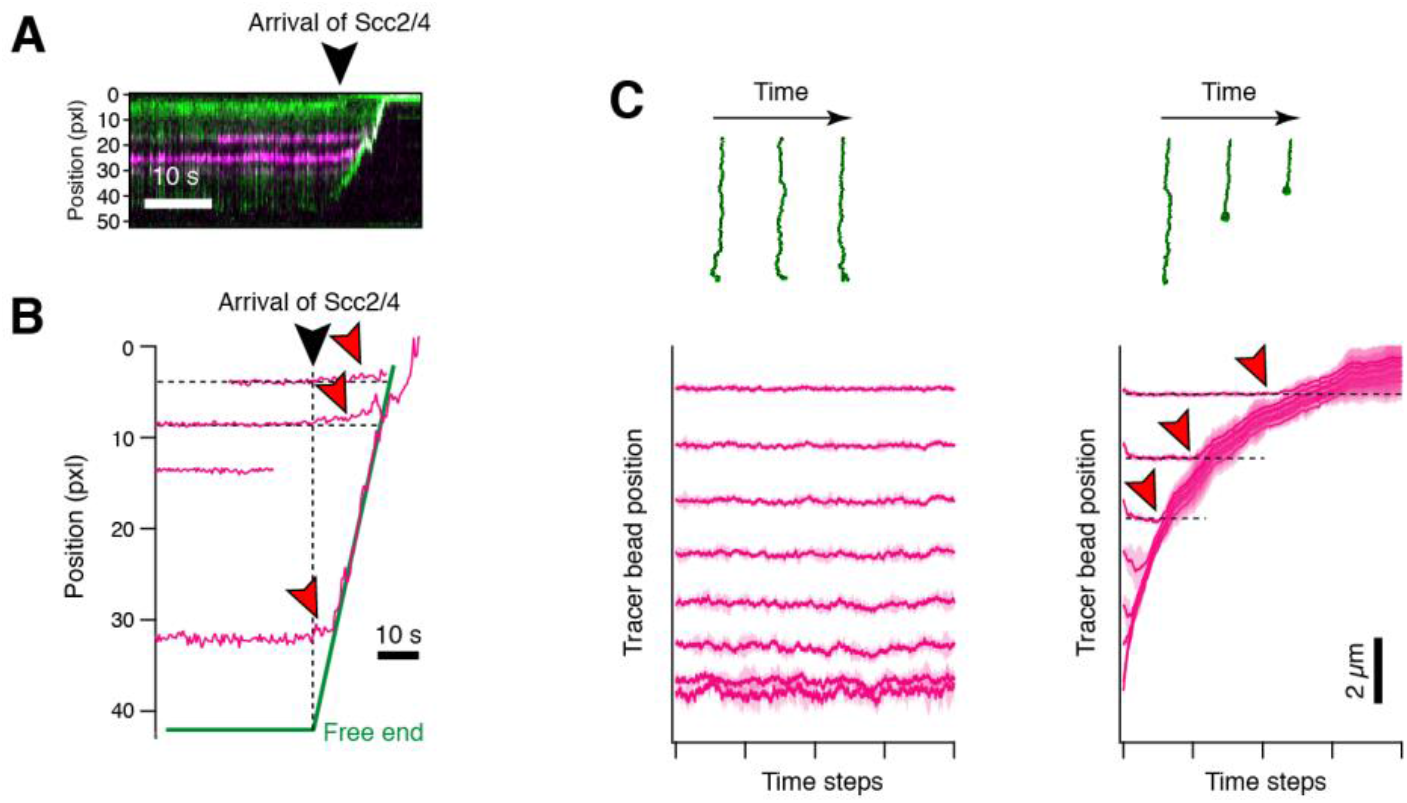
Internal DNA compaction. (**A**) Representative kymogram of Scc2/4-mediated DNA compaction with two EcoRI-E111Q tracer proteins (magenta) bound. (**B**) Tracked positions of tracer proteins upon Scc2/4 addition. The red arrows show movement of internal tracers distant from the condensing end. (**C**) Coarse-grained molecular dynamics simulations. Lambda DNA was represented with a Rouse model of 160 beads that were connected by segment lengths of ∼300 bp. Flow-stretching force was applied. Left: naked DNA, DNA is not compacted over time. Right: DNA with Scc2/4, which was modeled by introducing an attractive Lennard-Jones-like potential between the beads, DNA is compacted over time. Red arrows indicate the internal beads, which, unlike in experimental data (see (**C**)), do not move. Shaded areas indicate the standard deviation of triplicate simulation runs.

**Figure S5.**
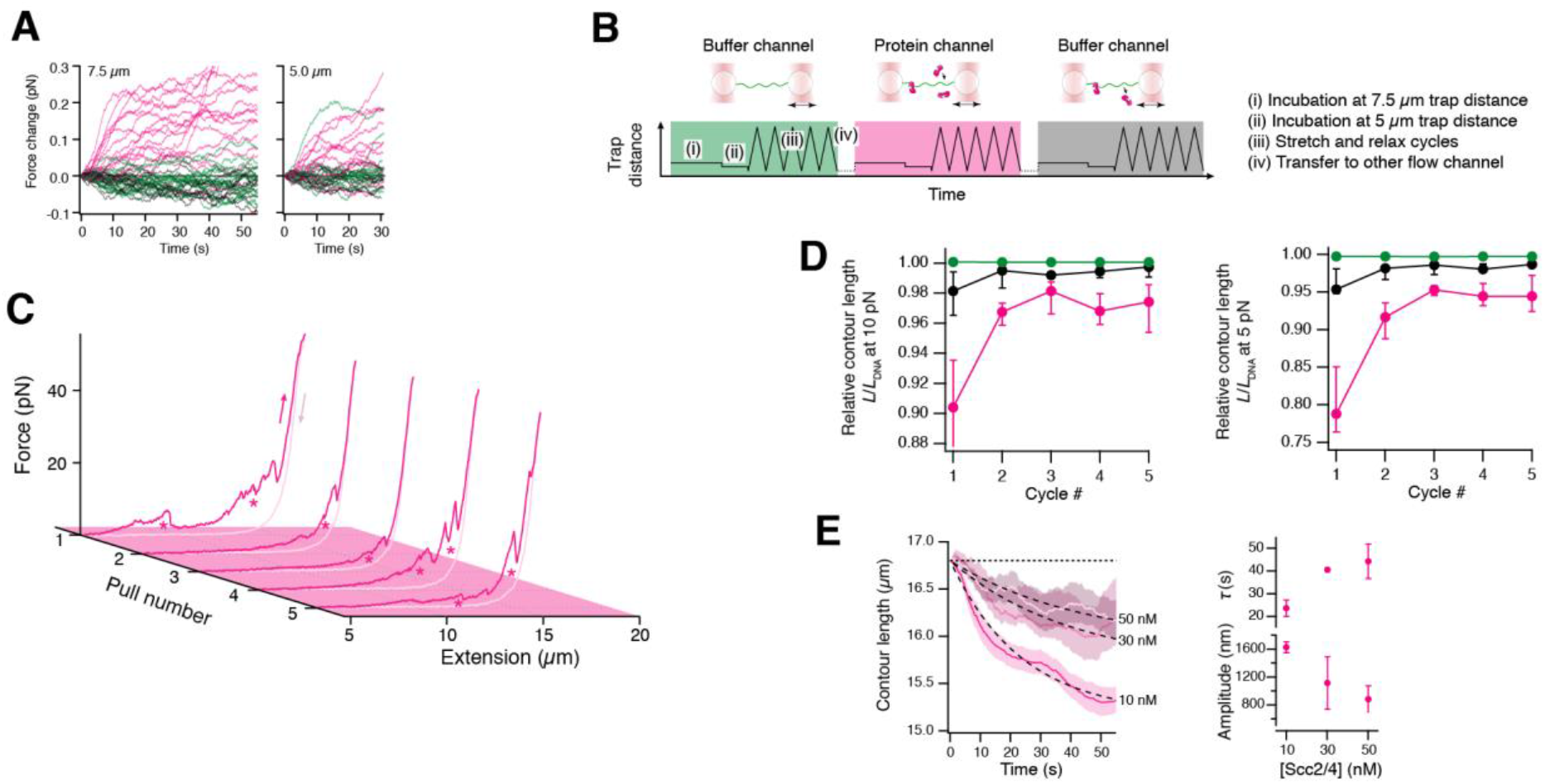
DNA compaction studied by optical tweezers. (**A**) Individual transient force responses of 15 lambda DNA molecules held at constant trap separation at the indicated initial forces. Green: without Scc2/4. Magenta: with 10 nM Scc2/4. Black: buffer wash after initial incubation with Scc2/4. Cf. **Fig. 5A**. (**B**) Experimental workflow. (**C**) Subsequent pulls of lambda DNA in a channel with 10 nM Scc2/4. The characteristic sawtooth pattern reforms in every cycle, observed compaction is marked by asterisks. (**D**) Relative compaction of tethers, as a function of pulling cycles. As loop formation is not instantaneous, the first cycle is significantly more compacted than the subsequent ones. (**E**) Time-dependent reduction of the tether contour length hold at 7.5 μm at different protein concentrations. Magenta traces represent averages. Dashed lines are fits to an exponential. Right: Amplitude and time constant of the exponential decrease in contour length.

**Figure S6.**
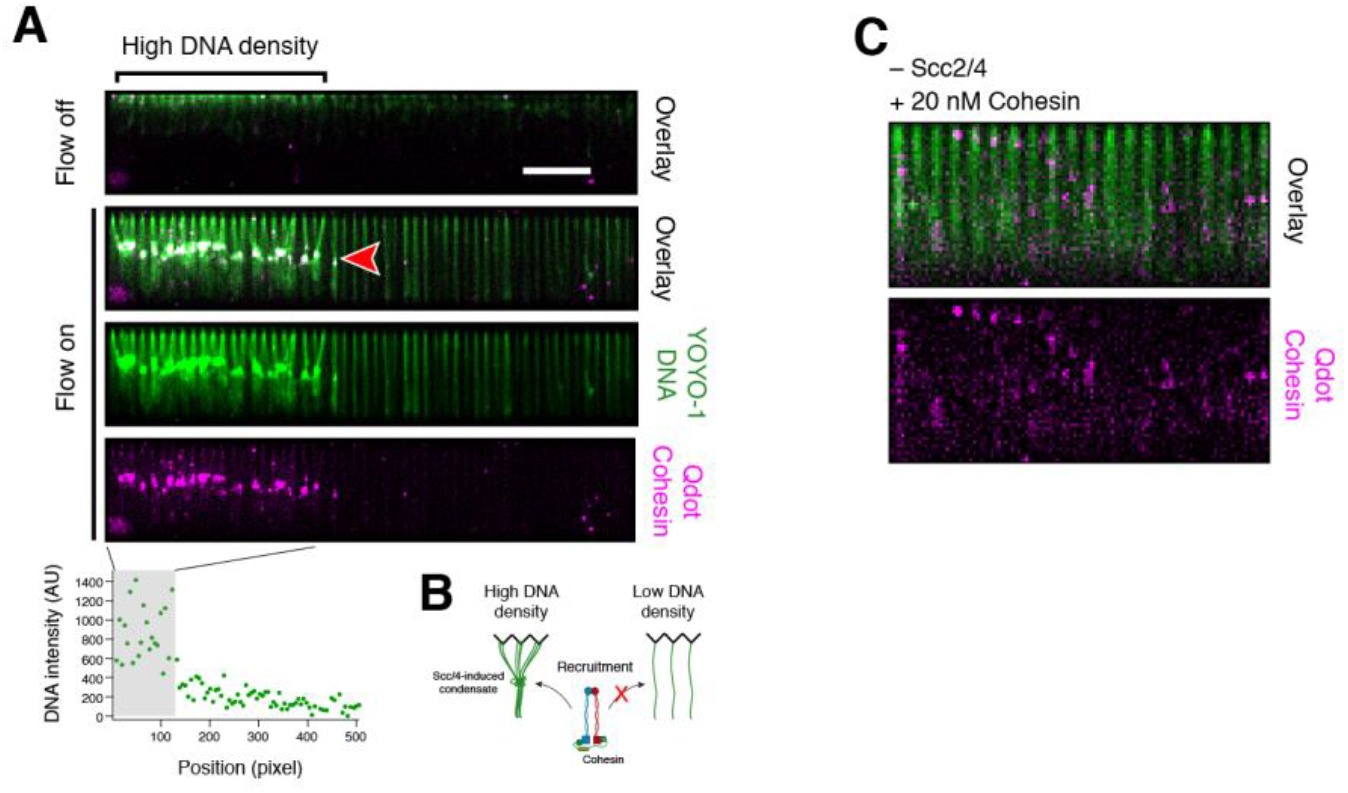
Cohesin binding behavior on DNA curtains. (**A**) 2 nM Qdot-labeled cohesin (magenta) and 8 nM Scc2/4 (dark) were applied simultaneously to single tethered DNA curtains (green) with a flow of 0.3 ml/min followed by incubation for 5 min without flow. Cohesin loading occurred primarily at dense DNA clusters created by Scc2/4-mediated condensation. DNA density was determined by fluorescence intensity. (**B**) Model for cohesin recruitment to Scc2/4-DNA co-condensates. (**C**) DNA curtain experiment without Scc2/4. 20 nM Qdot-labeled cohesin (magenta) were applied to single tethered DNA curtains (green) under continuous flow of 0.1 ml/min. Only few binding events occurred.

